# Allosteric deimmunisation of a *Salmonella* phosphatase enhances catalytic function

**DOI:** 10.1101/2025.11.17.688463

**Authors:** Soobon Ko, Sung-Hwan You, Yeojin Moon, Junsu Ko, Woong-Hee Shin, Yoonjoo Choi

**Affiliations:** Department of Biomedical Sciences, Chonnam National University Medical School, Hwasun-gun, Jeollanam-do, Republic of Korea; CNCure Biotech, Inc, Republic of Korea; Arontier Inc., Gangnam-Daero 241, Seocho-Gu, Seoul, Republic of Korea; Department of Biomedical Informatics, Korea University College of Medicine, Seoul, 02841, Republic of Korea; Department of Microbiology and Immunology, Chonnam National University Medical School, Hwasun-gun, Jeollanam-do, Republic of Korea; The National Immunotherapy Innovation Center, Hwasun-gun, Jeollanam-do, Republic of Korea

**Keywords:** Protein engineering, deimmunisation, allostery, bacteria cancer therapy, enzyme engineering

## Abstract

Genetically engineered *Salmonella* strains are promising vectors for cancer therapy, but their clinical use is limited by host immune responses. A key immunogenic antigen, the phosphatase PhoN, is metabolically essential for bacterial survival in the vitamin B6-depleted tumour microenvironment and thus cannot be simply deleted. While conventional deimmunisation targets surface residues, we present a strategy of allosteric deimmunisation by mutating non-surface, buried residues within the dominant h2 helix T-cell epitope. We hypothesised that the epitope is allosterically coupled to the distal active site gate, allowing for the simultaneous modulation of immunogenicity and enzymatic function. Using a computational pipeline, we designed deimmunised variants and used molecular dynamics (MD) simulations to predict their functional effects. The simulations revealed that mutations in the h2 helix allosterically controlled the gate’s flexibility. The enzymatic activities of the deimmunized variants were correlated with the MD predictions: two of the five designed variants exhibited 2∼3-fold increases in catalytic function. This work demonstrates that non-surface epitopes can be rationally engineered to not only ablate immunogenicity but also to allosterically enhance protein function.

## Introduction

Attenuated, tumour-homing bacteria such as *Salmonella* genus have emerged as promising agents for cancer therapy. These bacteria can selectively colonise and proliferate within the unique tumour microenvironment (TME) (*1*). Early clinical trials with engineered *Salmonella* strains have demonstrated safety (*2–7*), but a critical bottleneck remains, the host immune system (*8*). In particular, the adaptive response driven by CD4+ T cells rapidly clear the bacteria before they can fully exert their anti-cancer effects. This robust host immunity, while beneficial for clearing infections, poses a major barrier to therapeutic bacteria.

Protein immunogenicity represents a fundamental challenge in this context. Similar to biologic drugs that elicit anti-drug antibodies and neutralising immune responses (*9*), bacterial proteins can trigger strong T cell activation. Conventional deimmunisation strategies such as PEGylation physically mask epitopes but may compromise activity through steric interference (*10, 11*). Recent advances in computational deimmunisation by T-cell epitope removal have improved this process by jointly optimising immunogenicity and protein stability (*12–15*). However, these approaches generally confine mutations to solvent-exposed regions and overlook immunogenic motifs buried within structurally or functionally critical cores (*12, 16*).

Buried-site redesign remains exceptionally difficult, as internal residues are densely packed and energetically coupled to maintain protein integrity (*17–20*). Yet, it could decouple immune recognition from surface interactions and even uncover new routes for functional enhancement. We demonstrate this concept using *Salmonella* PhoN, a periplasmic acid phosphatase identified as one of the most immunodominant CD4+ T cell antigens across *Salmonella* serovars (*21*). While knocking out virulence genes is a common technique in such cases, PhoN is known indispensable for *Salmonella* survival in the vitamin B6(VB6)-depleted TME (*22*), catalysing the dephosphorylation of VB6 derivatives essential for bacterial metabolism (*23*). It thus represents both a key determinant of bacterial fitness and a major source of immunogenicity.

To resolve this paradox, we engineered PhoN variants that evade immune detection while enhancing enzymatic performance. Our approach departs from conventional surface-based designs by introducing targeted mutations within buried residues of the immunodominant region. The region is coupled to a gating loop that regulates substrate access to the active site (*24*). We hypothesised that subtle alterations in this buried region could both disrupt MHC II binding and fine-tune enzyme dynamics. Using a computational pipeline integrating epitope prediction, energy-based protein design, and molecular dynamics simulations, we generated PhoN variants predicted to eliminate known T cell epitopes while improving catalytic efficiency.

## Results

### Structural analysis of wildtype PhoN reveals a dynamic, ligand-sensitive active site gate

To establish a structural and dynamic baseline, we first analysed the wildtype (WT) *S. enterica* PhoN enzyme using the dimeric crystal structure (PDB ID: 2IPB). The immunodominant region (h2 helix, residues 49-62) is spatially distant from the enzyme’s active site, which is defined by key catalytic residues including R130, H156, and D191 (**Fig. 1A**). We hypothesised that these two distal regions, the T-cell epitope and the active site, are allosterically coupled via connecting helices (residues approximately 66-76) that form the gate to the substrate-binding pocket. To investigate the coupling hypothesis, we performed short all-atom molecular dynamics (MD) simulations (for 50 ns) using GROMACS (Ver. 2024.5) (*25*) with the AMBER99SB-ILDN force field (*26*).

**Figure 1.**
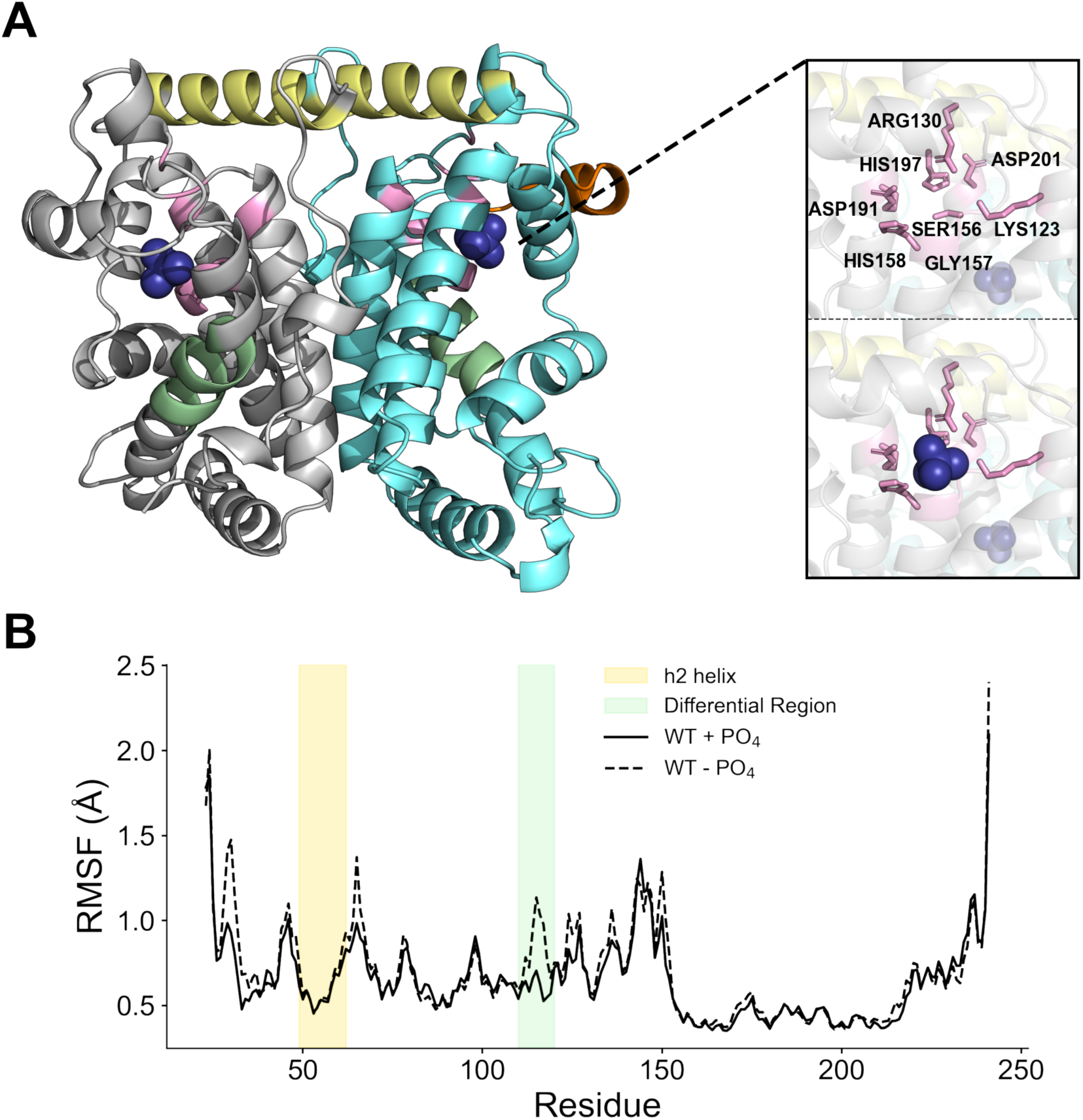
Structural dynamics of PhoN reveal ligand-dependent gating motion. **(A)** Dimeric structure (PDB ID: 2A96) of *Salmonella enterica* PhoN highlighting the h2 helix (residues 49–62) and the distal active site gate containing key catalytic residues (K123, R130, H156, G157, H158, D191, H197, and D201). **(B)** RMSF comparison between the phosphate-free (WT-PO_4_) and phosphate-bound (WT+PO_4_) states. Residues 110–120 exhibit elevated flexibility in the apo state, which becomes stabilised upon phosphate binding. The active-site gate and its motion are indicated in both structural and RMSF representations.

We compared the structural dynamics of the WT dimer in two distinct states: a ligand-free (*21*) state (WT-PO_4_) and a phosphate-bound (holo) state (WT+PO_4_). We first assessed the overall stability of both simulations by calculating the root mean square deviation (RMSD) relative to the starting crystal structure. Both systems reached equilibrium within the first 10 ns of the simulation and remained largely stable throughout the 50 ns production run (**Fig. S1**). We then analysed the per-residue flexibility by calculating the root mean square fluctuation (RMSF) for both simulations. This analysis revealed that though the two states are highly similar, certain local differences are observed between residues 110-120 (**Fig. 1B**). In the apo (WT-PO_4_) simulation, the helical region corresponding to the proposed active site gate (specifically residues 110-120) displayed significantly elevated flexibility. In contrast, this high degree of local motion was substantially dampened in the holo (WT+PO_4_) simulation upon the binding of the phosphate ligand.

This ligand-dependent change in mobility strongly supports the functional role of the local region (110–120) as a gate of the active site as depicted in **Fig. 1B**. The data suggests this gate is inherently mobile in the unbound state, presumably to facilitate substrate access to the catalytic pocket. Upon ligand binding, the gate becomes more ordered, likely to secure the substrate for the catalytic action of the active site residues. The demonstrated, innate flexibility of the gate region and its sensitivity to the ligand-bound state validate it as a key dynamic element, providing a clear biophysical rationale for our hypothesis that its motion could be allosterically modulated by targeted mutations within the coupled h2 helix.

### Computational non-surface residue engineering achieves successful deimmunisation

The target T-cell epitope in the h2 helix is composed of 20 amino acids in length (46 DDPAYRYDKEAYFKGYAIKG 65) (*21, 24*). Using a consensus of prediction tools (IEDB 2.22 (*27*), ProPred (*28*), and NetMHCIIpan-4.3 (*29*)), the 20-mer wildtype (WT) peptide was predicted to bind to multiple HLA class II alleles (**Fig. 2A**). In contrast to conventional deimmunisation approaches that target surface residues, we focused on non-surface residues within the immunodominant helix. Positions for mutagenesis were selected by filtering for residues with low solvent accessibility (**Fig. 2B**). We then employed the Rosetta design framework to generate stable, deimmunised sequences (*14, 30*). Among the total of 4,000 models generated, we selected five distinct variants (Var1 to 5) based on dual-objective criteria; 1) the lowest-energy conformations identified, and 2) the largest, most-converged sequence clusters. The resulting five variants, each possessing a unique combination of mutations within the non-surface h2 region (**Table 1**), were subsequently re-analysed in silico. This computational validation confirmed the success of our design protocol: all five variants were predicted to have no MHC-II binding events across all three prediction algorithms.

**Figure 2.**
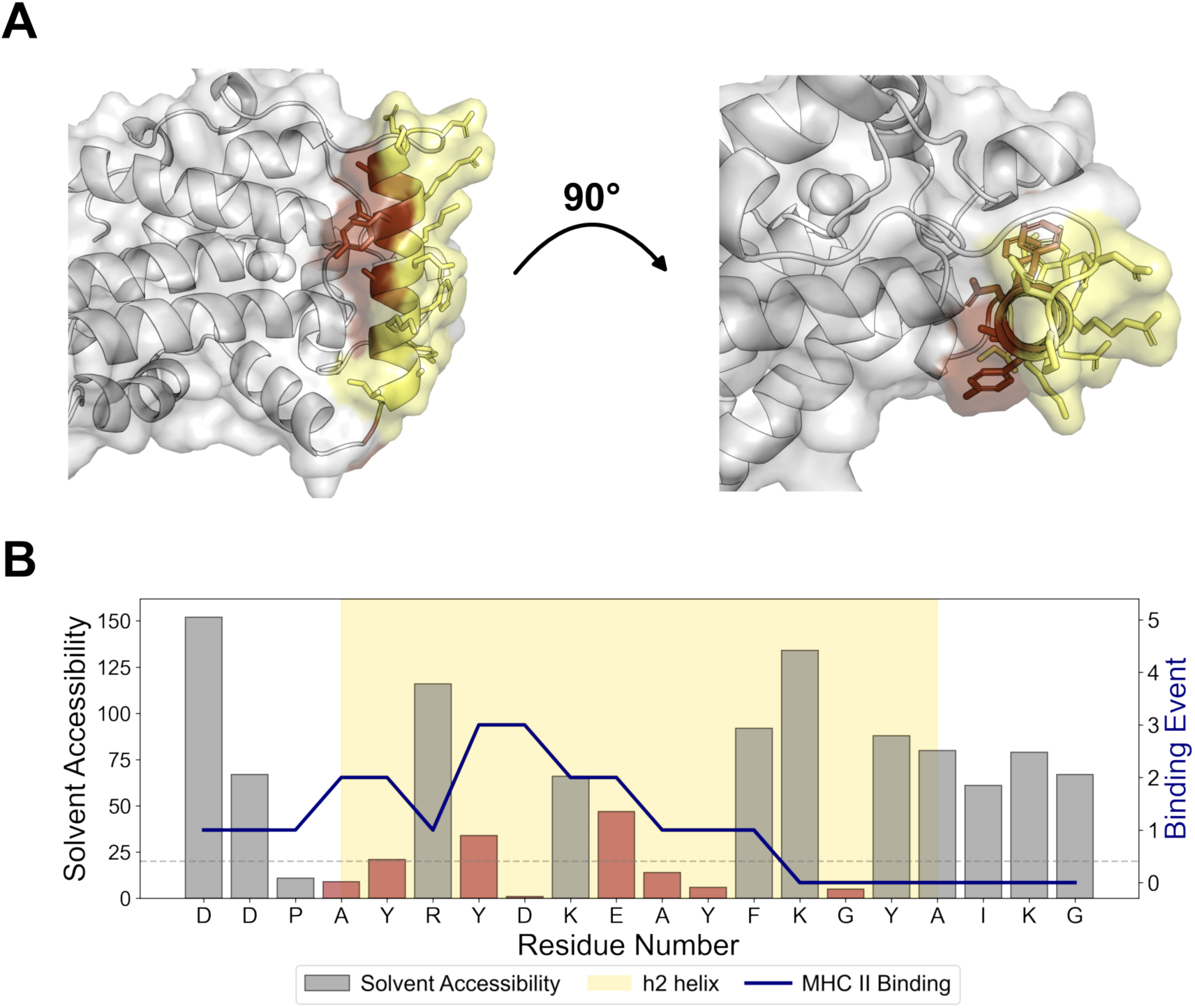
Structure-informed identification of buried immunogenic residues in the PhoN h2 helix. **(A)** Structural representation of the PhoN dimer showing the h2 helix (residues 49–62) in both side (left) and top (right) views. In the top-down view (right), the designed mutation sites (highlighted) are located in the buried interior of the h2 helix. **(B)** Predicted MHC-II epitope map overlaid with the solvent accessibility (ACC) profile. Epitope-prone regions with high predicted binding scores are indicated, while brown bars mark residues with low ACC values (< 50) targeted for mutation.

**Table 1.** Selected deimmunised PhoN variants and their predicted impact on HLA-DRB1*01:01 binding.

|  | Position |  |  |  |  |  |  |  |  | HLA-DRB1*01:01 |  |  |
| --- | --- | --- | --- | --- | --- | --- | --- | --- | --- | --- | --- | --- |
|  | 49 | 50 | 52 | 53 | 55 | 56 | 57 | 60 | 65 | NetMHCIIpan<br>4.3 | ProPred | IEDB |
| WT | A | Y | Y | D | E | A | Y | G | G | 1 | 4 | 4 |
| Var1 |  |  | A |  |  | H | A | E |  | 0 | 0 | 0 |
| Var2 | H | A | S | G |  | S | H | E |  | 0 | 0 | 0 |
| Var3 | E | D | E | G |  | C | H | E |  | 0 | 0 | 0 |
| Var4 |  | F | A |  | D | H | A | E |  | 0 | 0 | 0 |
| Var5 |  | F | A |  | D | H | A | E | D | 0 | 0 | 0 |

To experimentally validate these in silico predictions, we synthesised various forms of peptides corresponding to the T-cell epitope of the WT enzyme and the designed variants (**Table 2**). We then directly measured their binding affinities to HLA-DRB1*01:01. The results shows that the WT peptides exhibited significant binding as expected. In contrast, all other peptide variants corresponding to the variants yielded negligible binding events. These results confirms that that our computational redesign for deimmunisation successfully eliminated the dominant T-cell epitope.

**Table 2.** Experimental validation of deimmunised PhoN variants by MHC-II binding assay.

| Peptide ID | Label | Peptide Sequence | REVEAL<sup>®</sup> Score |
| --- | --- | --- | --- |
| 1 | Wildtype (46-65) | DDPAYRYDKEAYFKGYAIKG | 36.9 |
| 2 | Var 1 (46-65) | DDPAYRADKEHAFKEYAIKG | 0.1 |
| 3 | Var 2 (46-65) | DDPHARSGKESHFKEYAIKG | 0.1 |
| 4 | Var 3 (46-65) | DDPEDREGKECHFKEYAIKG | 0 |
| 5 | Var 4 (46-65) | DDPAFRADKDHAFKEYAIKG | 0.2 |
| 6 | Var 5 (46-65) | DDPAFRADKDHAFKEYAIKD | 0 |
| 7 | Wildtype (46-60) | DDPAYRYDKEAYFKG | 0.2 |
| 8 | Var 1 (46-60) | DDPAYRADKEHAFKE | 0.3 |
| 9 | Var 2 (46-60) | DDPHARSGKESHFKE | 0.1 |
| 10 | Var 3 (46-60) | DDPEDREGKECHFKE | 0.1 |
| 11 | Var 4 (46-60) | DDPAFRADKDHAFKE | 0 |
| 12 | Var 5 (46-60) | DDPAFRADKDHAFKE | 0.2 |
| 13 | Wildtype (51-65) | RYDKEAYFKGYAIKG | 39.1 |
| 14 | Var 1 (51-65) | RADKEHAFKEYAIKG | 0.3 |
| 15 | Var 2 (51-65) | RSGKESHFKEYAIKG | 0.1 |
| 16 | Var 3 (51-65) | REGKECHFKEYAIKG | 0.2 |
| 17 | Var 4 (51-65) | RADKDHAFKEYAIKG | 0.1 |
| 18 | Var 5 (51-65) | RADKDHAFKEYAIKD | 0 |

### Molecular dynamics simulations reveal allosteric control of the active site gate

To investigate the structural and functional consequences of our non-surface mutations, we performed 50 ns all-atom MD simulations for the WT enzyme and all five designed variants (Var1-5). We focused our analysis on the ligand-bound (holo, +PO_4_) state to specifically probe how the mutations influenced the enzyme’s dynamics in a catalytically relevant conformation. All six simulations demonstrated moderately stable RMSD values throughout the 50 ns trajectories (**Fig. S2**). However, RMSF analysis revealed significant and divergent allosteric effects.

The mutations introduced in the non-surface residues in h2 helix induced pronounced dynamic changes at the spatially distant active site gate (**Fig. 3A**). This effect was particularly evident in variants Var2 and Var3, which both exhibited a dramatic increase in flexibility within this specific gate region (**Fig. 3B**). In fact, our simulation of the Var2+PO_4_ complex, the phosphate ligand was unable to be secured by the destabilised gate, and thus was subsequently expelled from the binding pocket (**Video S1**). This observation led to the speculation that variants Var2 would be catalytically impaired or inactive.

**Figure 3.**
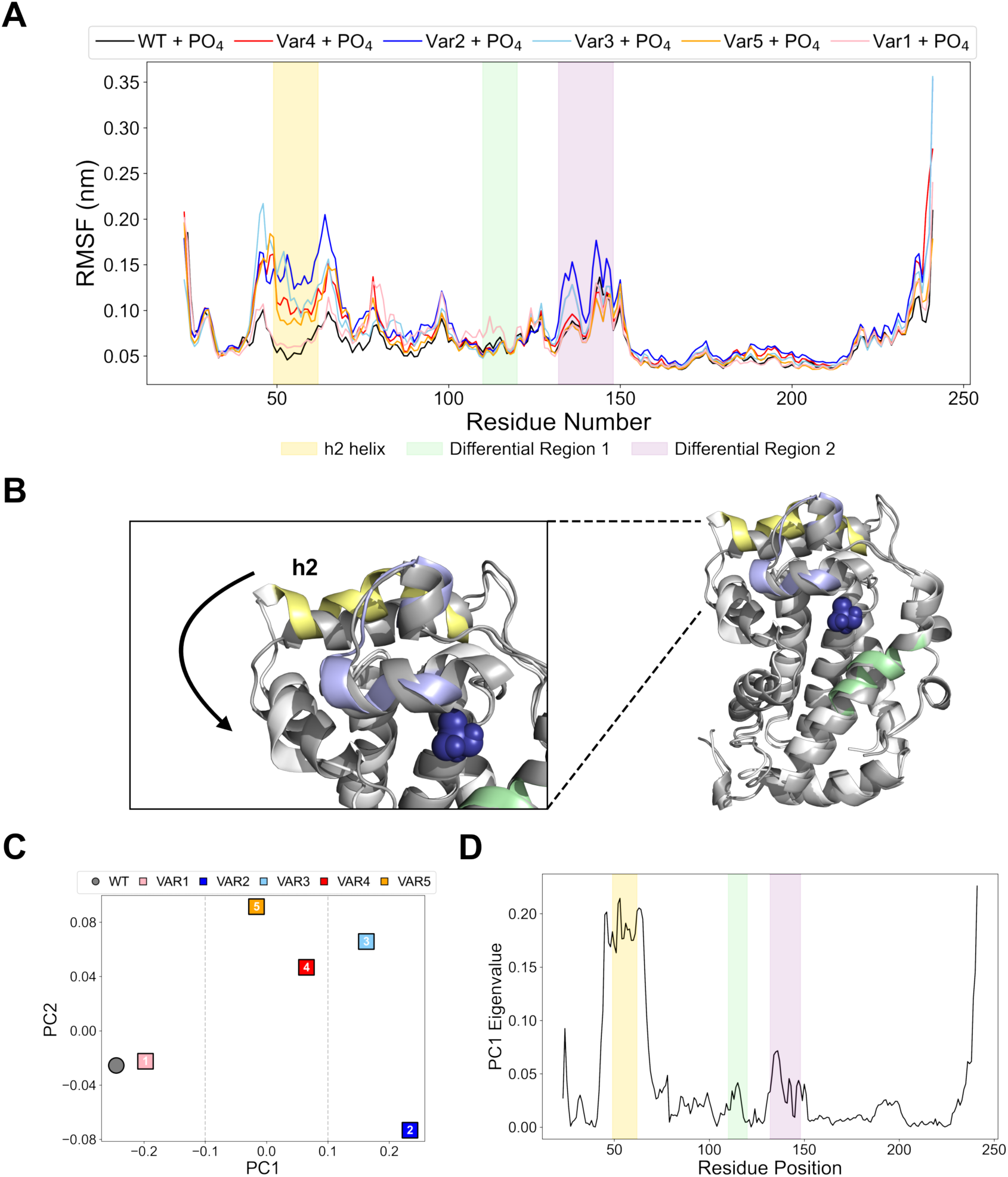
Variant-induced modulation of local and global dynamics in PhoN. **(A)** RMSF profiles of the PO_4_-bound wildtype and five designed variants. Regions with divergent fluctuation patterns among variants are highlighted in light purple. **(B)** Structural comparison between WT and Var2, where mutations in the h2 helix induce local perturbations in the adjacent α-helix. **(C)** PCA of RMSF profiles for six trajectories (WT and variants), revealing clear separation of variant groups along PC1 according to their flexibility patterns. **(D)** Residue-wise eigenvalue profile for PC1 represents the contribution of each residue to the principal component. Peaks correspond to residues with the largest influence on PC1-associated motions.

To classify the collective dynamic signatures of all variants, we performed a principal component analysis (PCA) on the RMSF values. The resulting plot of the first two principal components demonstrates that the variants may roughly segregate into three distinct dynamic ensembles (**Fig. 3C**): (WT/Var1), (Var2/Var3), and (Var4/Var5). We further analysed the principal components to physically interpret this dynamic separation.

An analysis of the eigenvectors for the first principal component (PC1) revealed that the residues with the highest eigenvector weights were located, except for the h2 helix region, almost exclusively in the active site gate (**Fig. 3D**). This confirms that the primary dynamic difference captured by the PCA is precisely the allosterically-controlled flexibility of this functional gate, rather than nonspecific motion elsewhere in the protein. This computational segregation allowed us to hypothesise that these three dynamic ensembles, defined by the specific mobility of the active site gate, would correspond directly to three distinct functional outcomes: (i) baseline activity (WT/Var1), (ii) impaired activity (Var2 and possibly Var3), and (iii) a potentially other altered activity (Var4/Var5).

### In vitro enzymatic assays validate the computationally-predicted functional classes

To experimentally test the functional hypotheses generated from the MD simulations, we proceeded with the heterologous expression and purification of all the variants. The variants were successfully expressed in *E. coli* BL21 (DE3) and purified from the soluble lysate fraction using Ni-NTA affinity chromatography (**Fig. S3**). The enzymatic function of these purified proteins was then quantified using an in vitro acid phosphatase assay, which measures the hydrolysis of the substrate p-nitrophenyl phosphate (pNPP).

The experimental results provided a direct and striking validation of the three distinct functional ensembles predicted by our in silico dynamic analysis (**Fig. 4**). First, Var1, which was found in the WT-like dynamic ensemble, exhibited a phosphatase activity functionally indistinguishable from the WT baseline as shown in the MD simulations. The MD simulation of Var2+PO_4_ showed potential catalytic impairment and Var3 was clustered with Var2 according to the PCA analysis. The enzyme activity results show that the two indeed showed approximately 50% activity reduction compared to the WT. The reduced functionality is in agreement with the MD simulation data for this cluster, which demonstrated excessive, high-amplitude motion in the active site gate. The activity loss provides a strong experimental validation for this computationally-observed failure mechanism, which confirms that excessive gate flexibility is detrimental to enzyme function.

**Figure 4.**
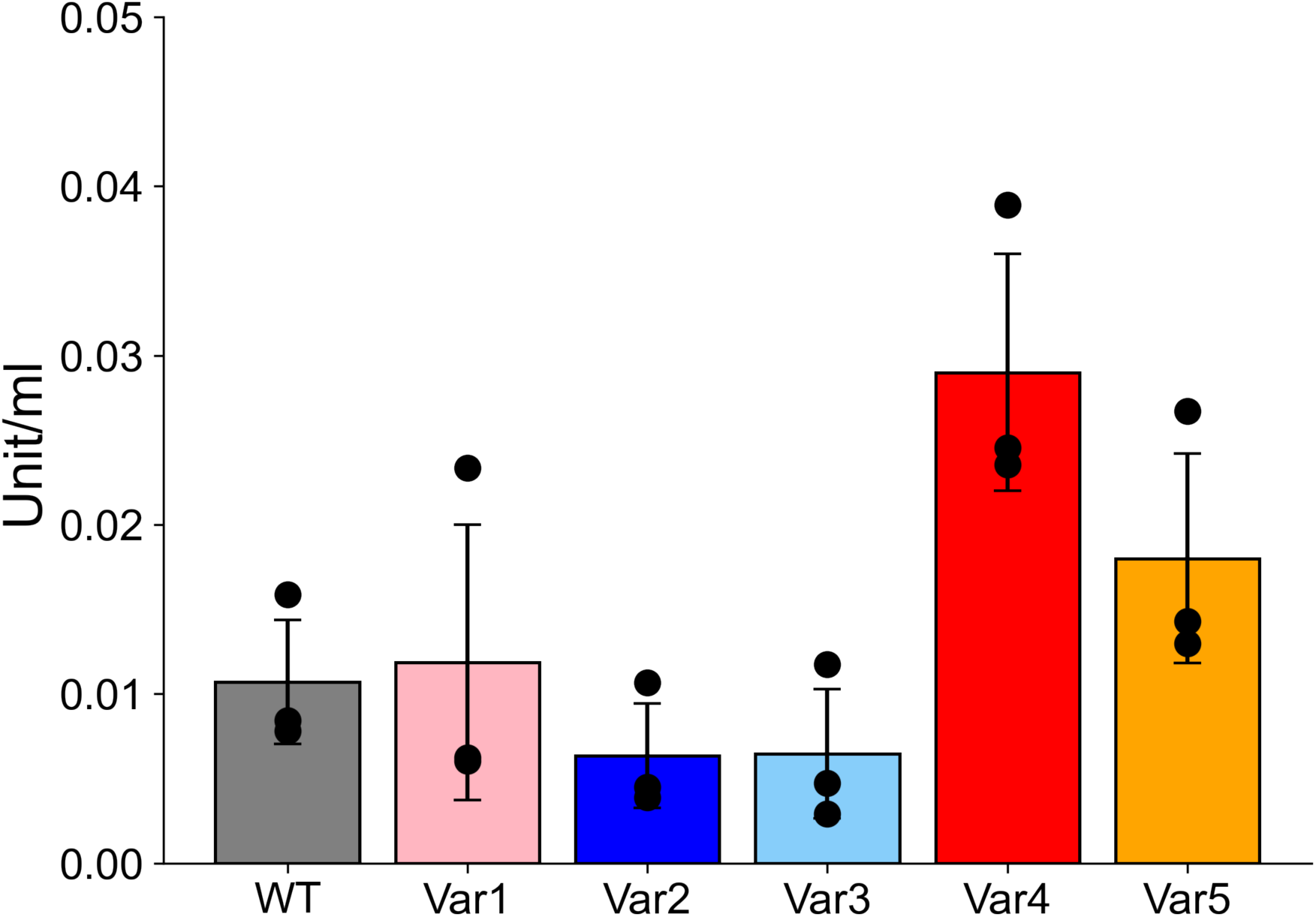
Acid phosphatase activity of wildtype and deimmunised PhoN variants. Enzymatic activity of purified PhoN proteins measured using the p-nitrophenyl phosphate (pNPP) assay. Bars represent mean standard deviation from three independent experiments (n = 3). Variant activities are normalised to the wildtype value.

On the other hand, Var4 and Var5, which the PCA segregated into another dynamic ensemble were both experimentally confirmed as highly active functional variants. Both variants yielded approximately 2∼3-fold increases in activity. The in vitro validation of this third functional class confirms that the unique dynamic signature corresponds to an optimised, supra-physiological function.

The direct, one-to-one correspondence between the three computationally predicted dynamic clusters and the corresponding experimentally measured functional outcomes is a powerful validation of our hypothesis. The results confirm that the non-surface h2 helix functions as an allosteric control element for the distal active site gate. Furthermore, it demonstrates that the computational design strategy, which targets non-surface residues and thus make changes in the dynamics of the functional site, can be used not only to ablate immunogenicity but also to tune enzyme dynamics to achieve a significant, multi-fold enhancement of catalytic function.

## Discussion

Therapeutics from non-human sources present practical difficulties such as the immunogenicity problem. The host adaptive immune response to foreign proteins is an inevitable trade-off that can limit therapeutic efficacy (*31*). We faced this challenge with the *Salmonella* phosphatase PhoN, which is both a dominant CD4+ T cell antigen and a metabolically essential enzyme for bacterial survival in the vitamin B6-depleted tumour microenvironment (*21–23*). While adopting computational protein deimmunisation techniques is expected to reduce the immunogenicity problem for an independent use of the protein, there is a risk in applying conventional methods for deimmunisation in bacteria-based therapy. In general, deimmunising mutations are intentionally introduced in surface residues and regions distant from active sites to minimise the risk of destabilisation. However, targeting surface residues of a protein component of therapeutic agents may cause unexpected detrimental alterations in protein-protein interactions.

Here, we show that functionally enhanced deimmunisation can be achieved by directly targeting non-surface residues and controlling the dynamics of the active site. The conventional goal of protein deimmunisation typically aims to preserve baseline function. However, our strategy delivers enzyme variants catalytically superior to its wildtype progenitor. This engineered PhoN represents an ideal candidate for next-generation *Salmonella* vectors that can evade human immune detection while more efficiently executing its essential metabolic function within the TME.

The mechanistic basis for this functional tuning was established by the strong correlation between computational simulations and experimental assays. The molecular dynamics simulations revealed that the mutations in the h2 helix function as an allosteric control element. There appears to be an optimal dynamic balance for the active site gate. Excessive flexibility is detrimental, while the wildtype state (WT/Var1) is stable but sub-optimal. The dynamic ensemble occupied by Var4 and Var5 represents an optimised dynamic state that results in an approximately 4-fold increase in catalysis.

This work also challenges the conventional wisdom of protein redesign for deimmunisation. Typically, deimmunising mutations are confined to solvent-exposed surface residues to avoid disrupting the structural integrity. Mutations in buried regions are known to be far more sensitive and destabilising. Our strategy, however, was predicated on the hypothesis that this particular buried region was not just an immunogenic liability but a functional opportunity. By targeting these non-surface residues, we demonstrated that the introduction of deimmunising mutations can be the very mechanism used to rationally re-tune the enzyme’s dynamic landscape. This concept of allosteric deimmunisation, where the epitope is the key for functional control, may prove to be a powerful strategy for engineering other therapeutic proteins where epitopes are found in structurally or allosterically significant regions.

While our results present a significant advance at the molecular level, this study also has limitations that define the scope for future work. Our MHC-II binding assay was limited to the reported single allele (HLA-DRB1*01:01). For practical applications, it should be expanded to a broader panel of common HLA alleles. This study was conducted entirely at the protein level. Future studies will therefore focus on translating these molecular findings into a therapeutic context. The definitive test will be to engineer attenuated *Salmonella* strains that express the successful variant gene. By introducing these strains into tumour models, we can directly test the therapeutic hypothesis: that an enhanced B6-salvage capability will lead to superior bacterial proliferation and persistence in the TME, while the deimmunised profile will delay clearance by the host adaptive immune system.

## Methods

### Computational Methods

#### T-cell Epitope Prediction

Three computational tools were used to predict T-cell epitopes: NetMHCIIpan 4.3 (*29*), ProPred (*28*), and the IEDB recommended method (version 2.22) (*27*). Each tool was utilized to predict MHC Class II peptide binding across multiple human leukocyte antigen (HLA) alleles. A sliding window approach was applied across all prediction methods, generating overlapping peptides of varying lengths, 15mers for NetMHCIIpan and the IEDB recommended method, and 9mers for ProPred. The IEDB recommended method and NetMHCIIpan analysed 27 HLA Class II alleles (*32*), with peptides ranked within the top 5% percentile classified as binders (10% for the IEDB method). ProPred predicted epitope binding against eight representative HLA-DRB1 alleles (DRB1*01:01, 03:01, 04:01, 07:01, 08:01, 11:01, 13:01, and 15:01). A 5% percentile threshold was applied for ProPred to classify peptides as potential binders. For all prediction methods, epitope scores were computed as the linear sum of binding events across all analysed alleles.

#### Computational Design of PhoN Variants

The PhoN crystal structure (PDB ID: 2IPB) was retrieved from the Protein Data Bank and prepared for computational design using Rosetta (Ver. 3.13). Residues within the predicted T-cell epitope regions were selected as potentially designable positions.

This set was further refined based on two key filters, solvent accessibility and evolutionary conservation.

First, to target non-surface residues, solvent exposure was quantified using the DSSP algorithm (*33*). Residues with ACC < 50 were classified as having reduced solvent exposure and considered for mutation. Second, to ensure that proposed mutations would maintain protein functional conservation, an evolutionary constraint analysis was performed. Homologous protein sequences were retrieved via protein BLAST (*34*) searches against the refseq database (maximum 5,000 sequences). A multiple sequence alignment (MSA) was then generated using Muscle (Version 3.8.31) (*35, 36*) to establish natural amino acid variation at each site. Only mutations observed in naturally occurring homologs and also predicted to reduce epitope binding scores were prioritized.

Using this filtered set of non-surface, evolutionarily-vetted positions, amino acid substitutions were designed using Rosetta’s MHCEpitopeEnergy framework (*30*). Each chain was designed independently, generating 2,000 sequence variants per chain. Final designs were selected based on high sequence convergence across multiple simulations, with the lowest-energy variant from the converged sequence group selected for further experimental characterization.

#### Molecular Dynamics Simulation and Analysis

All-atom molecular dynamics (MD) simulations were performed using GROMACS 2024 (*25, 37*) with the AMBER99SB-ILDN force field (*26*). Initial dimeric structures of PhoN variants, both with and without bound PO_4_ ligands, were solvated in a dodecahedral box with a minimum 1.0 nm buffer to the protein, using the TIP3p water model. The system was neutralized, and 0.15 M NaCl was added to mimic physiological ionic strength.

Energy minimization was carried out using the steepest descent algorithm. Equilibration followed in two phases: 100 ps under NVT conditions using the V-rescale thermostat, followed by 200 ps of NPT pre-equilibration using the Berendsen barostat, and 500 ps under NPT conditions using the Parrinello–Rahman barostat. Production runs were conducted for 50 ns with a 2 fs timestep at 310 K and 1 bar. All bond lengths were constrained using the LINCS algorithm, and long-range electrostatics were handled with the PME method. Analyses of root-mean-square deviation (RMSD), root-mean-square fluctuation (RMSF) were performed using GROMACS built-in utilities to evaluate structural stability, local flexibility changes, and collective dynamic motions, particularly in the active-site gate region.

### Experimental Methods

#### Protein expression and purification

The PhoN genes (wildtype and variants 1–5) were cloned into the pET21-a vector using NdeI and XhoI restriction sites, ensuring in-frame fusion with a C-terminal 6×His tag. The recombinant pET21-a constructs were transformed into BL21 (DE3) *Escherichia coli* (New England Biolabs) and selected using ampicillin. Transformed BL21 (DE3) colonies were first cultured overnight in Luria-Bertani (LB) medium supplemented with ampicillin at 37°C for 16 hours. The overnight culture was then diluted 1:100 into fresh LB medium (10 mL + ampicillin) and incubated at 37°C with shaking (220 rpm) until the optical density at 600 nm (OD600) reached 0.8-1.0. The culture was then cooled to 16°C for 30 minutes, followed by induction with 0.5 mM isopropyl β-D-1-thiogalactopyranoside (IPTG) at 16°C for 16 hours.

Induced cells were harvested by centrifugation at 15,000g for 10 minutes and the resulting pellet was resuspended in lysis buffer (phosphate-buffered saline, pH 7.4, supplemented with 1x protease inhibitor cocktail (Thermo Fisher Scientific) and 10 mM imidazole). Cells were lysed by sonication on ice at 25% amplitude (Utlrasonic processor Z412619, Sigma-Aldrich), using a pulse duration of 2s and a pulse off-time of 2s, for a total sonication time of 20s. The lysate was then centrifuged at 15,000g for 1 minute at 4°C, and the supernatant was subjected to Ni-NTA affinity purification using a Ni-NTA spin kit (Qiagen) according to the manufacturer’s protocol. The purified proteins were eluted with PBS (pH 7.4) containing 500 mM imidazole.

#### MHC Class II Binding Assay

The ProImmune REVEAL® MHC Class II Binding Assay was employed to assess the MHC-binding affinity of 18 peptides, including both the wildtype sequence and selected Rosetta-designed variants, for the HLA allele complex DRA*01:01/ DRB1*01:01. In this assay, antibody-labelled peptides emit a detectable signal upon binding to MHC Class II molecules, thereby confirming both binding and the stability of the resulting MHC-peptide complex. Measurements were taken at two distinct time points (0 and 24 hours), and each peptide was then scored for binding to an HLA molecule on a scale of 0 to 100 relative to a high-affinity positive control baseline provided by ProImmune. Peptides with binding scores exceeding 15% of the positive control were considered to exhibit significant binding. All peptides were synthesized using Prospector PEPscreen® technology, and purity was verified via MALDI-TOF mass spectrometry.

#### Acid phosphatase assay

The enzymatic activity of purified PhoN proteins was assessed using the Acid Phosphatase Assay Kit (Sigma-Aldrich) following the manufacturer’s instructions. Briefly, 50 μg of purified protein was mixed with the p-nitrophenyl phosphate (pNPP) substrate solution and incubated at 37°C for 5-10 minutes. The reaction was terminated by adding 0.5 N NaOH (stop solution), and absorbance was measured at 405 nm using a spectrophotometer (Infinite 200 PRO, Tecan). A positive control with 0.05 μmole/mL p-nitrophenol (pNP) was used as standard solution to relatively quantify the phosphatase activities of samples. The specific activity was expressed as units/mL, where one unit of acid phosphatase will hydrolyze 1 μmole of pNPP per min at pH 4.8 at 37°C.

## Acknowledgments

Korea Health Technology R&D Project through the Korea Health Industry Development Institute (KHIDI), funded by the Ministry of Health & Welfare, Republic of Korea (grant number: RS-2025-25459531). The National Research Foundation of Korea (NRF) grant funded by the Korean government (MSIT) under the program Medical Research Center for Innovative Control of Cardiovascular Remodeling Diseases (RS-2025-02213506) to YC. Bio & Medical Technology Development Program of NRF funded by MSIT (NRF-2022M3E5F3081268 and NRF-2024M3A9J4006525) to JK. Bio & Medical Technology Development Program of NRF (NRF-2022M3E5F3081268) to WS. SK is funded by RS-2024-00462620.

