## Supplementary material for "Allosteric deimmunisation of a *Salmonella* phosphatase enhances catalytic function": Fig. S1

### Supplementary Figures

**Figure S1. Structural stability of the PhoN dimer under $PO_4$-bound and unbound conditions.** Root-mean-square deviation (RMSD) plots of the PhoN dimer during 50 ns molecular dynamics simulations performed with (violet) and without (black) $PO_4$ ligands.

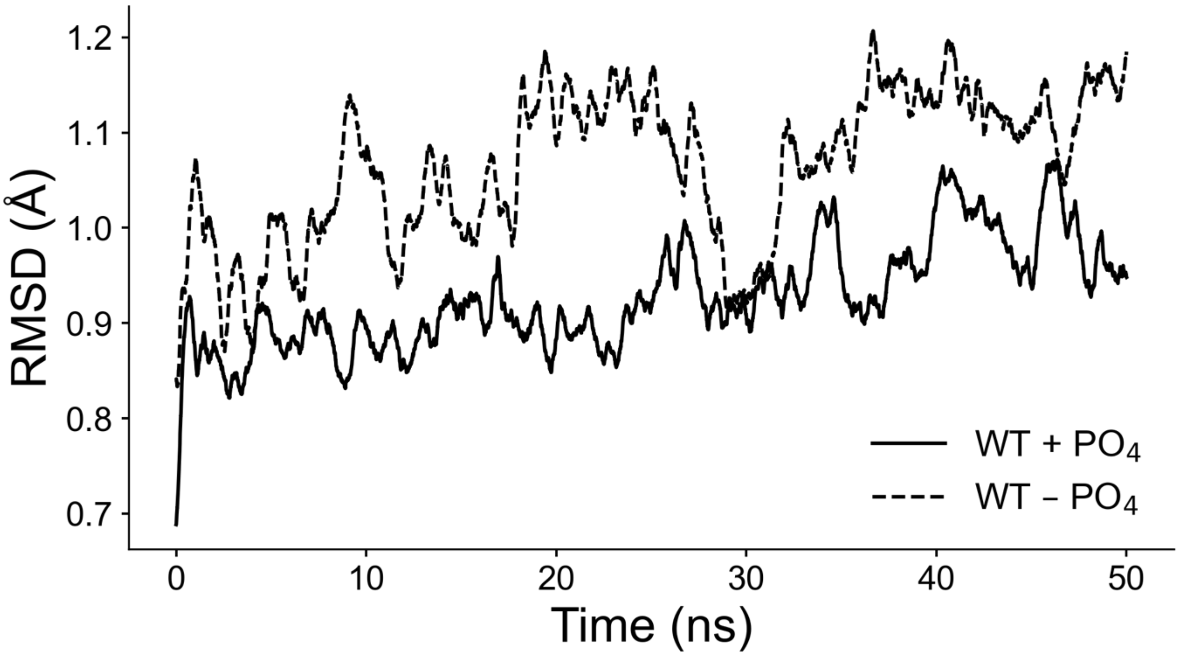

**Figure S2. Structural stability of wildtype and variants during $PO_4$-bound molecular dynamics simulations.** Root-mean-square deviation (RMSD) trajectories of the wild-type and five designed variants (Var1-5) over 50 ns under $PO_4$-bound conditions. All six simulations showed stable RMSD convergence, indicating that the introduced mutations did not compromise the overall structural integrity of PhoN.

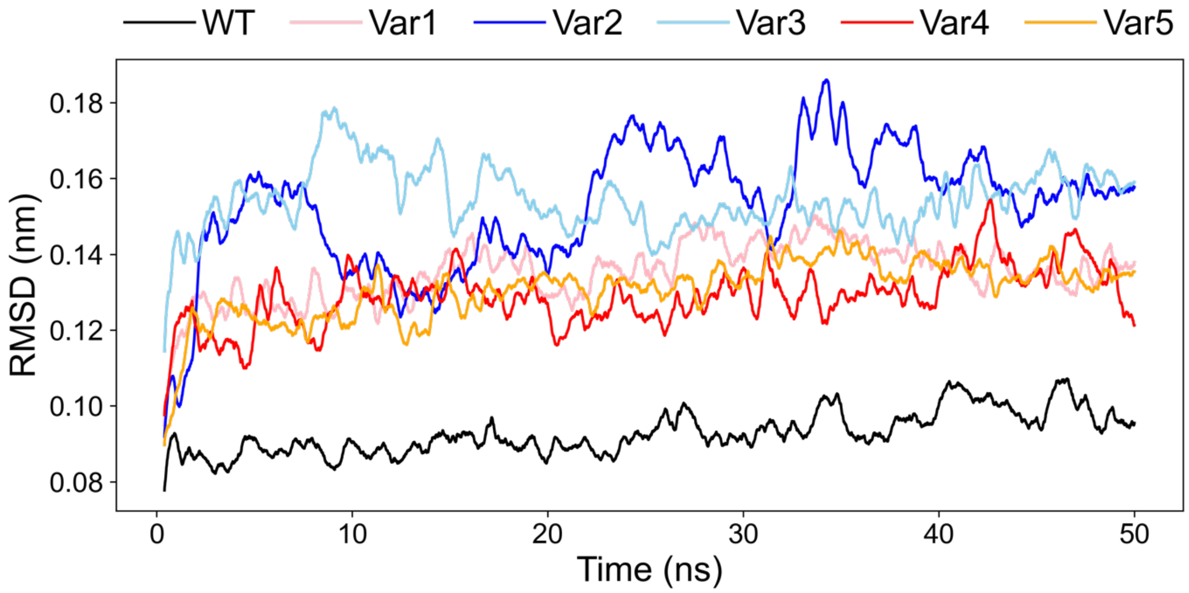

**Figure S3. Expression and purification of wildtype and deimmunised PhoN variants.** SDS-PAGE gel showing the soluble expression and Ni–NTA affinity purification of the wild-type (WT) and designed variants (Var1-5) expressed in *E. coli* BL21 (DE3). All proteins were successfully purified from the soluble fraction and migrated near the expected molecular weight (~27 kDa).

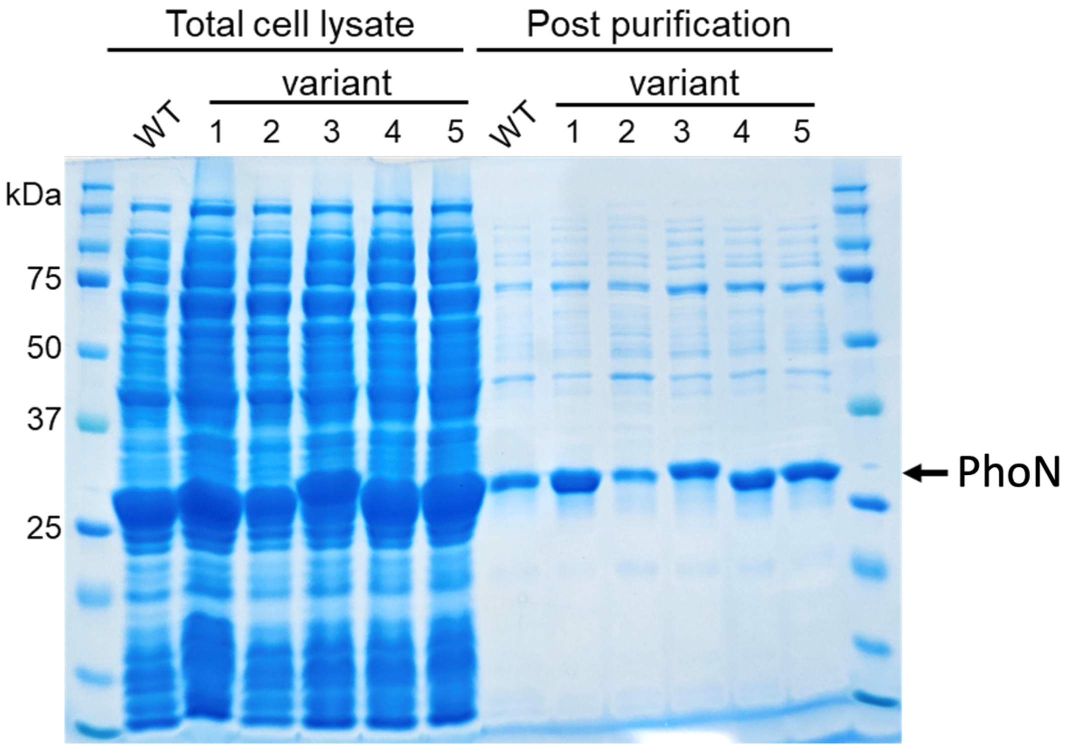
